# HSPA8-mediated stability of the CLPP protein regulates mitochondrial autophagy in cisplatin-resistant ovarian cancer cells

**DOI:** 10.1101/2023.08.24.554577

**Authors:** Xinxin Kou, Xiaoxia Yang, Zheng Zhao, Lei Li

**Affiliations:** Department of Gynecology, Cancer Hospital Affiliated to Zhengzhou University, Zhengzhou, China

**Keywords:** Cisplatin resistance, ovarian cancer, caseinolytic protease P, heat shock protein family A member 8, mitophagy

## Abstract

**Background:** Currently, platinum agents remain the mainstay of chemotherapy for ovarian cancer (OC). However, cisplatin (DDP) resistance is a major reason for chemotherapy failure. Thus, it is extremely important to elucidate the mechanism of resistance to DDP.

**Methods:** We establish 2 DDP-resistant ovarian cancer cell lines and find that caseinolytic protease P (CLPP) is significantly downregulated in the DDP-resistant cell lines when compared to wild-type ovarian cancer cell lines (SK-OV-3 and OVcar3). Next, we investigate the functions of CLPP in the DDP-resistant and wild-type ovarian cancer cells using various assays including cell counting kit-8 assays, western blotting, immunofluorescent staining, and reactive oxygen species (ROS) and apoptosis detection.

**Results:** Our experiments show that CLPP knockdown significantly increase the half maximal inhibitory concentration (IC_50_) and mitophagy of wild-type SK-OV-3 and OVcar3 cells, while CLPP overexpression reduces the IC_50_ values and mitophagy of DDP-resistant SK-OV-3 and OVcar3 cells. Next, we perform database predictions and experiments to show that heat shock protein family A member 8 (HSPA8) regulates CLPP protein stability. The dynamic effects of the HSPA8/CLPP axis in the ovarian cancer cells were also examined. HSPA8 increases mitophagy and the IC_50_ values of SK-OV-3 and OVcar3 cells, but inhibits their ROS production and apoptosis. In addition, CLPP partly reverses the effects induced by HSPA8 in the SK-OV-3 and OVcar3 cells.

**Conclusions:** CLPP increases the DDP resistance of ovarian cancer by inhibiting mitophagy and promoting cellular stress. Meanwhile, HSPA8 promotes the degradation of CLPP protein by inducing its stability.

## Introduction

Ovarian cancer is the 3rd most common gynecologic malignancy worldwide, and the most lethal of the gynecologic malignancies [1]. Currently, the incidence and mortality rates of ovarian cancer continue to increase [2]. The mainstream chemotherapy for epithelial ovarian cancer (EOC) is a platinum agent combined with taxane [3]. However, patients eventually develop tolerance to cisplatin (DDP)-based therapy after a few cycles of treatment, which further increases the mortality rate [4]. Thus, there is an urgent need to elucidate the underlying mechanism of DDP resistance in ovarian cancer.

DDP acts by binding to DNA and subsequently causing DNA damage, inhibition of DNA replication, and an induction of cell apoptosis [5]. Based on DDP pharmacological studies and its biological activities, various mechanisms may contribute to DDP resistance, including an increased cellular efflux of drugs [6], reduced drug influx [7], inhibition of apoptosis [8], dysregulation of DNA damage repair systems [9], enhanced activity of drug-metabolizing enzymes [10, 11], and an altered cellular microenvironment [12-14]. The reasons for DDP resistance in DDP-resistant cancer cells include a reduction in drug accumulation, DDP inactivation resulting from its reaction with glutathione and metallothionein, and a rapid repair of DNA lesions [9, 15]. Copper efflux transporters, such as Cu-transporting P-type ATPases (ATP7A and ATP7B), have been shown to regulate the efflux of DDP [16, 17]. Recent studies have shown that changes in mitochondrial function abnormal autophagy play key roles in DDP resistance in ovarian cancer [18-20], which suggests that mitochondrial molecules are involved in DDP resistance.

Mitochondrial caseinolytic protease P (CLPP) is a serine protease located in the mitochondrion matrix and is involved in mitochondrial protein metabolism (proteostasis) and oxidative stress by facilitating the degradation of misfolded or damaged proteins, and thus the maintenance of protein metabolism homeostasis [21, 22]. Due to the functions of CLPP in mitochondria, it has multiple effects on tumors [23]. In acute myeloid leukemia, the genetic and chemical activation of CLPP selectively kills cancer cells by degrading respiratory chain protein substrates and disrupting mitochondrial structure and function; however, CLPP has no effects on non-cancerous cells [24]. In pancreatic ductal adenocarcinoma (PDAC), the chemical activation of CLPP increases the degradation of respiratory chain complexes, which causes an endoplasmic reticulum stress response that suppresses the growth of PDAC cells [25]. In glioblastoma, CLPP activators induce synthetic lethality by inhibiting oxidative energy metabolism and reducing cell viability [26]. In EOC, the mitochondrial deficits induced by CLPP inhibit the growth and metastasis of EOC cells [27]. However, the role of CLPP in DDP resistance in ovarian cancer remains unclear.

Heat shock protein family A member 8 (HSPA8) belongs to the heat shock protein 70 (HSP70) family [28]. It plays important physiological roles in protein metabolism and homeostasis, which both depend on constant protein degradation and resynthesis [29]. Under normal or stressful conditions, eukaryotic cells remove misfolded proteins via autophagy. In chaperone-mediated autophagy, HSPA8 recognizes and targets cytosolic proteins with a signature exposed pentapeptide motif (KFERQ) [30]. After being recognized by HSPA8 and binding to lysosomal-associated membrane protein 2A, the target proteins are translocated into the lysosomal lumen for degradation [31].

In this study, we explored the effects of CLPP on mitophagy in wild-type or DDP-resistant ovarian cancer cells. Specifically, our database predictions and experiments indicated that the *HSPA8* gene might regulate the stability of CLPP. Taken together, our findings suggest that HSPA8-mediated stability of the CLPP protein regulates mitochondrial autophagy in DDP-resistant ovarian cancer cells. Our results elucidate one of the mechanisms by which drug resistance occurs, provide new insights into cisplatin resistance in ovarian cancer, and will facilitate the development of new clinical therapies.

## Methods

### Cell lines and cell transfection

Human ovarian cancer cell lines SK-OV-3 (ATCC^®^ HTB-77^TM^) and OVcar3 (ATCC^®^ HTB-161^TM^) were purchased from the American Type Culture Collection (ATCC) (Manassas, VA, USA) and cultured in Roswell Park Memorial Institute Medium 1640 (Gibco, Waltham, MA, USA) supplemented with 10% fetal bovine serum (Gibco) at 37°C in a 5% CO_2_ atmosphere. DDP-resistant SK-OV-3 and OVcar3 cells were developed from the wild-type SK-OV-3 and OVcar3 cells by using a stepwise procedure in which the cells were exposed to increasing concentrations of DDP in culture medium for > 6 months.

### Real-time PCR analysis

TRIzol reagent (Invitrogen, Carlsbad, CA, USA) was used to extract the total RNA from cells and tissue samples. A reverse transcription kit (Takara, Tokyo, Japan) was used as per the kit’s instructions to perform reverse transcription. The quantitative polymerase chain reaction was performed by using SYBR Green Master Mix on a LightCycler 480 PCR system (Roche Diagnostics, Indianapolis, IN, USA). Glyceraldehyde-3-phosphate dehydrogenase served as an internal control. Relative levels of gene expression were calculated using the 2^−ΔΔ^^Ct^method, and glyceraldehyde-3-phosphate dehydrogenase served as an internal control. The primer sequences used for RT-PCR are shown in Table S1.

### Immunoblotting

The total proteins were extracted from treated cells, and the protein concentration in each extract was determined using the bicinchoninic acid protein assay (Pierce, Rockford, IL, USA). Next, a 40 μg aliquot of protein from each extract was separated by Tris-glycine sodium dodecyl-sulfate polyacrylamide gel electrophoresis (4% to 20%) and the protein bands were transferred onto polyvinylidene-fluoride membranes (Sigma-Aldrich, St. Louis, MO, USA), which were subsequently blocked with non-fat milk. The membranes were then incubated for 8–10 h with the following primary antibodies: anti-CLPP (ab124822, Abcam, Cambridge, UK), anti-PTEN-induced kinase 1 (PINK1) (ab300623, Abcam), anti-parkin (ab77924, Abcam), anti-(HSPA8) (ab265664, Abcam) anti-Microtubule-associated protein light chain 3 (LC3B) (ab48394, Abcam), and anti-β-actin antibody (ab8227, Abcam). The primary antibody was then detected after incubation with a secondary antibody conjugated with horseradish peroxidase. The protein signals were visualized with an enhanced chemiluminescence kit (Pierce).

### Cell counting kit-8 (CCK-8) assays

Cell viability was determined by performing CCK-8 assays on transfected and/or treated cels. After replacing the used media with fresh media, the cells were transferred into 96-well plates. Next, 20 μL of CCK-8 solution was added to each well and the cells were incubated for 3 hours. Finally, the optical density (OD) of each well was measured with a microplate reader at 450 nm.

### IF staining

Ovarian cancer cells were fixed for 30 min with 4% polyformaldehyde and then permeabilized for 10 min with 0.6% Triton X-100 before being blocked with goat serum. Next, following an overnight incubation at 4□with anti-CLPP (ab124822, Abcam) or anti-LC3B (ab48394, Abcam), the cells were incubated for 1 h in the dark with a secondary antibody conjugated to fluorescein isothiocyanate (FITC; Invitrogen). Next, the cell nuclei were stained with 4’,6-diamidino-2-phenylindole staining solution (ab104139, Abcam) at an ambient temperature, and the cells were imaged with a laser microscope (Olympus, Japan).

### ROS detection

The treated cells were harvested and incubated with dichloro-dihydro-fluorescein diacetate (DCFH-DA) (10 μM) for 30 min in the dark at 37□. Following incubation, the cells were harvested with trypsin/ethylenediaminetetraacetic acid and their intracellular levels of reactive oxygen species (ROS) were measured by flow cytometry as previously described [32].

### Cell apoptosis detection

Apoptotic cells were detected using an Annexin V-FITC Apoptosis Detection Kit (Keygen, China). In brief, the cancer cells were harvested with 0.25% pancreatin; after which, they were washed twice with phosphate buffered saline, and resuspended in 500 μL of binding buffer. The cells were then incubated with 5 μL of antibody against Annexin V-FITC and 5 μL of propidium iodide for 15–20 min in the dark. After incubation, the apoptotic cells were detected with a BD Accuri C6 flow cytometer (Becton Dickinson, Franklin Lakes, NJ, USA).

### Statistical analysis

All data were analyzed using IBM SPSS Statistics for Windows, Version 20 software and results are expressed as a mean value ± standard deviation. GraphPad 7.0 software (GraphPad Software, Inc., USA) and Biorender (Biorender-JavaShuo) was used to draw graphs. The Student’s *t*-test was used to compare mean values between 2 groups, and one-way analysis of variance was used to analyze differences among multiple groups.

## Results

### CLPP expression was downregulated in the DDP-resistant ovarian cancer cells

First, we established the DDP □resistant cell lines (SK-OV-3/DDP and OVcar3/DDP) by gradually the increasing the cisplatin concentration in SK-OV-3 and OVcar3 cells. We then verified the viability of the SK-OV-3 and OVcar3 cells under both normal conditions and DDP treatment conditions. As shown in *Figure 1A*, constant exposure to DDP significantly increased the survival rates of the SK-OV-3 and OVcar3 cells. The IC_50_ values of the DDP-resistant SK-OV-3/DDP cells and OVcar3/DDP cells were significantly increased when compared to those of the untreated wild-type SK-OV-3 and OVcar3 cells, which indicated successful establishment of the DDP □resistant ovarian cancer cell lines. After establishing the DDP□resistant cell lines, qRT-PCR and immunoblotting studies showed that the levels of CLPP messenger RNA (mRNA) and CLPP protein in the cisplatin-resistant groups were lower than those in the cisplatin-sensitive groups (*Figures 1B-1C*). Immunofluorescence (IF) staining also showed reduced CLPP protein levels in the SK-OV-3/DDP and OVcar3/DDP cells (Figure 1D). The results demonstrated that CLPP expression was significantly decreased in the DDP-resistant ovarian cancer cells when compared to the wild-type SK-OV-3 and OVcar3 cells.

**Figure 1.**
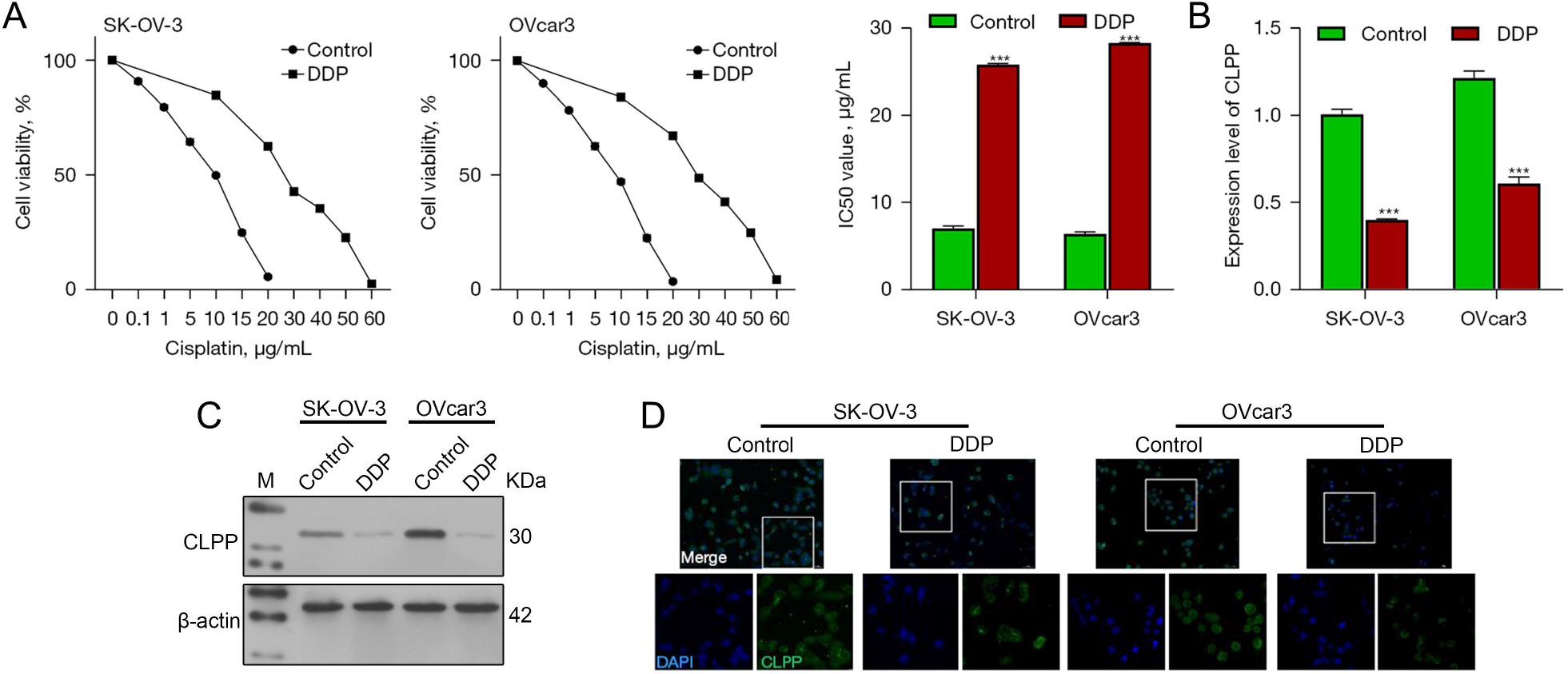
CLPP expression was upregulated in DDP-resistant ovarian cancer cells. (A) SK-OV-3 and OVcar3 cells were treated with DDP (0, 0.1, 1.0, 5.0, 10.0, 15.0, 20.0, 30.0, 40.0, 50.0, or 60.0 μg/mL) and examined for cell viability by the CCK-8 assay. DDP-resistant ovarian cancer cells were established. (B) The levels of CLPP mRNA in wild-type and DDP-resistant ovarian cancer cells were examined by real-time PCR. (C) The levels of CLPP protein in the cells were examined by immunoblotting. (D) The distribution of CLPP protein in the cells was examined by immunofluorescent staining (magnification x200 and x400). ***, P < 0.001, compared to the control group. CLPP, caseinolytic protease P; DDP, cisplatin; CCK-8, cell counting kit-8; PCR, polymerase chain reaction. Control, wild type SK-OV-3, and OVcar3 cell lines.

### Effects of CLPP knockdown in wild-type ovarian cancer cells

As shown in Figure 2A, CLPP knockdown significantly increased the IC_50_ values of the SK-OV-3 and OVcar3 cells. Studies have shown that the autophagy process of cancer cells is involved in resistance to chemotherapy [33]. Therefore, we next examined the levels of mitochondrial autophagy-related proteins (i.e., PINK1and Parkin) as well as the levels of autophagy associated proteins (LC3II/I) in cells being treated with DDP (IC_25_). CLPP interference was successfully achieved in the wild-type OVcar3 and SK-OV-3 cell lines (*Figure 2B,2C*). The SK-OV-3 and OVcar3 cells with CLPP knockdown were exposed to different concentrations of DDP (0–20 μg/mL), and IC_50_ values were calculated based on cell viability. Immunoblotting results showed that CLPP knockdown increased the levels of PINK1, Parkin, and LC3-II proteins (*Figure 2C*). The levels of LC3B protein in the SK-OV-3 and OVcar3 cells were also increased by CLPP knockdown (*Figure 2D*). These results indicated that CLPP knockdown can promote autophagy and mitochondrial autophagy in cancer cells. To further explore the oxidative stress and apoptosis of the cells, the ROS levels and apoptotic rates were detected by flow cytometry. As shown in *Figure 2E,2F*, CLPP knockdown decreased the ROS levels and apoptosis rates of wild-type SK-OV-3 and OVcar3 cells being treated with DDP (IC_25_).

**Figure 2.**
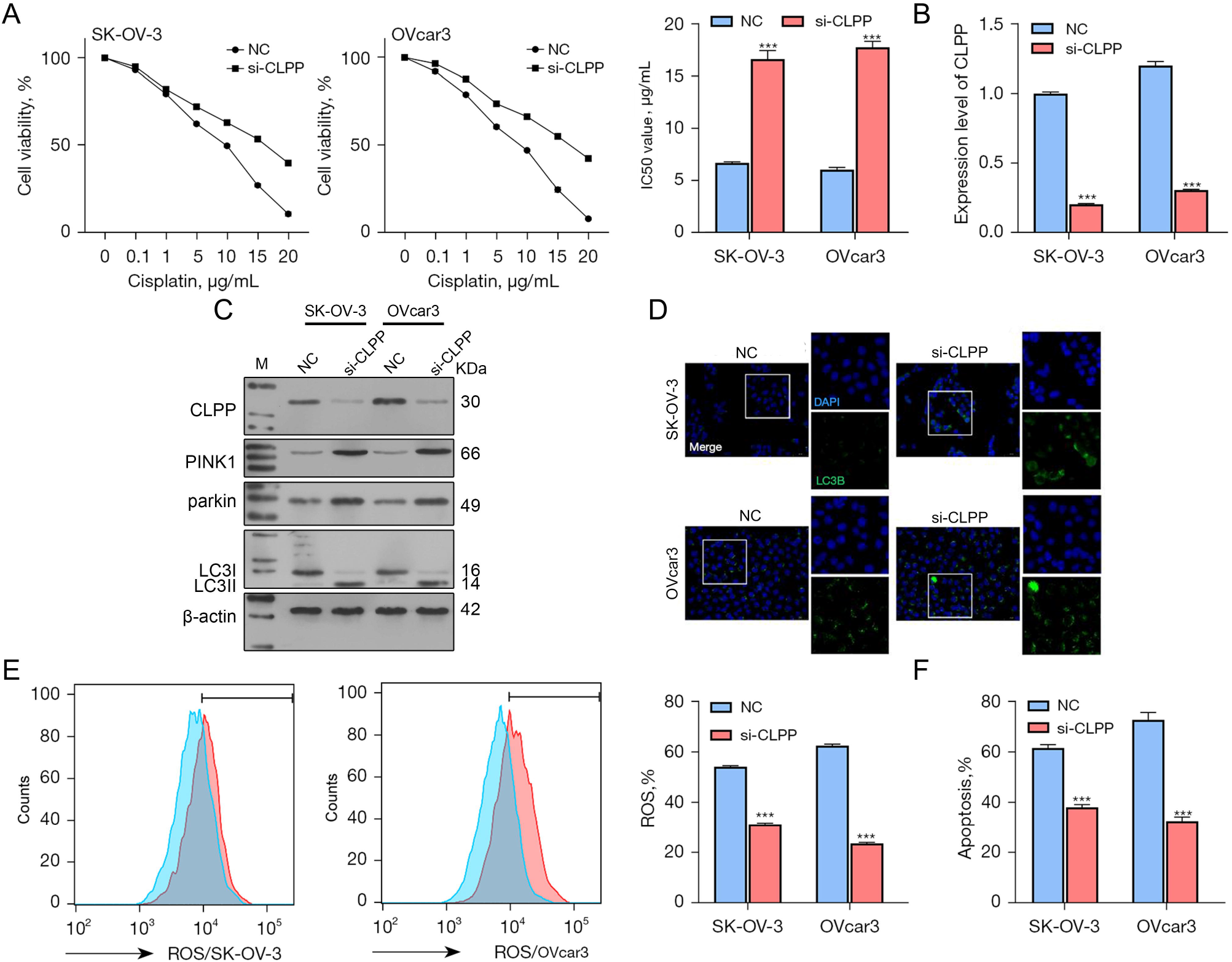
The effects of CLPP knockdown on wild-type ovarian cancer cells. (A) Wild-type ovarian cancer cells were transfected with a CLPP interference sequence and then treated with different concentrations of DDP; the IC_50_ values were calculated based on cell viability. (B) The levels of CLPP mRNA in the cells were examined by real-time PCR. (C) The levels of CLPP, PINK1, Parkin, and LC3II/I proteins were examined by immunoblotting. (D) The distribution of LC3B protein in the cells was examined by immunofluorescent staining (magnification x200 and x400). (E) ROS levels were measured by flow cytometry. (F) Cell apoptosis was examined by flow cytometry. ***, P < 0.001, compared to the NC group. NC, negative control; CLPP, caseinolytic protease P; DDP, cisplatin; ROS, reactive oxygen species; si, small interfering; PCR, polymerase chain reaction.

### Effects of CLPP overexpression in the DDP-resistant ovarian cancer cells

After determining the effects of CLPP in the wild-type ovarian cancer cells, we subsequently investigated the effects of CLPP in cisplatin-resistant ovarian cancer cells. As shown in Figure 3A CLPP overexpression significantly decreased the IC_50_ values of the DDP-resistant SK-OV-3 and OVcar3 cells. Our data also showed that CLPP overexpression was successfully achieved in the DDP-resistant SK-OV-3 and OVcar3 cell lines (*Figure 3B,3C*). Next, the DDP-resistant SK-OV-3 and OVcar3 cells with CLPP overexpression were exposed to different concentrations of DDP (0– 60 μg/mL), and IC_50_ values were calculated based on cell viability. Immunoblotting results showed that CLPP overexpression decreased the levels of PINK1, Parkin, and LC3II/I protein expression in cells the being treated with DDP (IC_25_) (*Figure 3C*). The levels of LC3B protein were also decreased in the DDP-resistant SK-OV-3 and OVcar3 cells with CLPP overexpression (*Figure 3D*). CLPP overexpression increased the ROS levels and apoptosis rates in the DDP-resistant SK-OV-3 and OVcar3 cells undergoing DDP treatment (IC_25_) (*Figure 3E,3F*). Furthermore, out data showed that overexpression of CLPP could reduce the levels of cellular and mitochondrial autophagy in cisplatin-resistant cells, increase the ROS levels and apoptosis rates of ovarian cancer cells, and thus improve the drug sensitivity of ovarian cancer cells.

**Figure 3.**
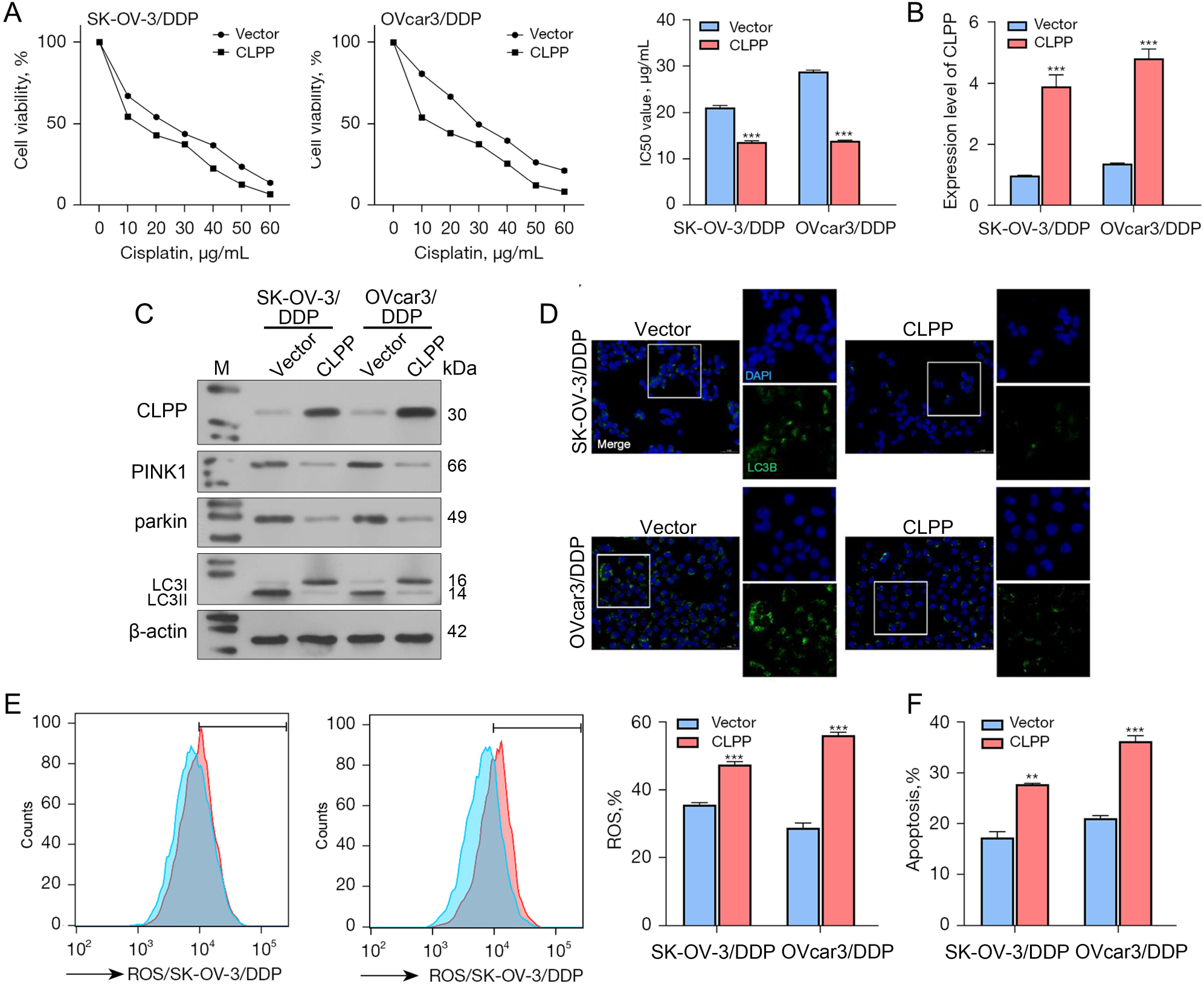
The effects of CLPP overexpression on DDP-resistant ovarian cancer cells. (A) DDP-resistant ovarian cancer cells were transfected with CLPP overexpression plasmids and subsequently treated with different concentrations of DDP; the IC_50_ values were calculated based on cell viability. (B) The levels of CLPP mRNA in the cells were examined by real-time PCR. (C) The levels of CLPP, PINK1, Parkin, and LC3II/I proteins were examined by immunoblotting. (D) The distribution of LC3B protein in the cells was examined by immunofluorescent staining (magnification x200 and x400). (E) ROS levels were measured by flow cytometry. (F) Cell apoptosis was examined by flow cytometry. **, P < 0.01; ***, P < 0.001, compared to the vector group. Vector, Empty carrier; CLPP, caseinolytic protease P; DDP, cisplatin; ROS, reactive oxygen species; si, small interfering; PCR, polymerase chain reaction.

### HSPA8 affected CLPP protein stability in ovarian cancer cells

After revealing that CLPP was involved in the DDP resistance of ovarian cancer, we investigated the underlying mechanism for its effect on DDP resistance. Our initial database predictions and experiments suggested that HSPA8 was involved in regulating CLPP protein stability. Thus, to explore the interaction between HSPA8 and CLPP, HSPA8 overexpression was achieved in the wild-type SK-OV-3 and OVcar3 cell lines (*Figure 4A*). Our results showed that CLPP mRNA levels remained unchanged after HSPA8 overexpression (*Figure 4B*). Conversely, CLPP protein levels were decreased by HSPA8 overexpression (*Figure 4C*). Similarly, HSPA8 knockdown was performed in the DDP-resistant SK-OV-3 and OVcar3 cell lines (*Figure 4D*). The levels of CLPP mRNA were not altered by HSPA8 knockdown (*Figure 4E*), while the CLPP protein levels were increased by HSPA8 knockdown (*Figure 4F*). Next, cycloheximide (CHX) was used to inhibit total cellular protein synthesis. Under conditions of CHX treatment, HSPA8 overexpression promoted the degradation of CLPP protein in SK-OV-3 and OVcar3 cells (*Figure 4G*). To further verify this process, a proteasome inhibitor (MG132) was used to treat HSPA8-overexpressing cells. Subsequent results showed that MG132 reversed the reductions in CLPP expression induced by HSPA8 (*Figure 4H*). These results indicated that HSPA8 reduces CLPP protein level by facilitating its stability.

**Figure 4.**
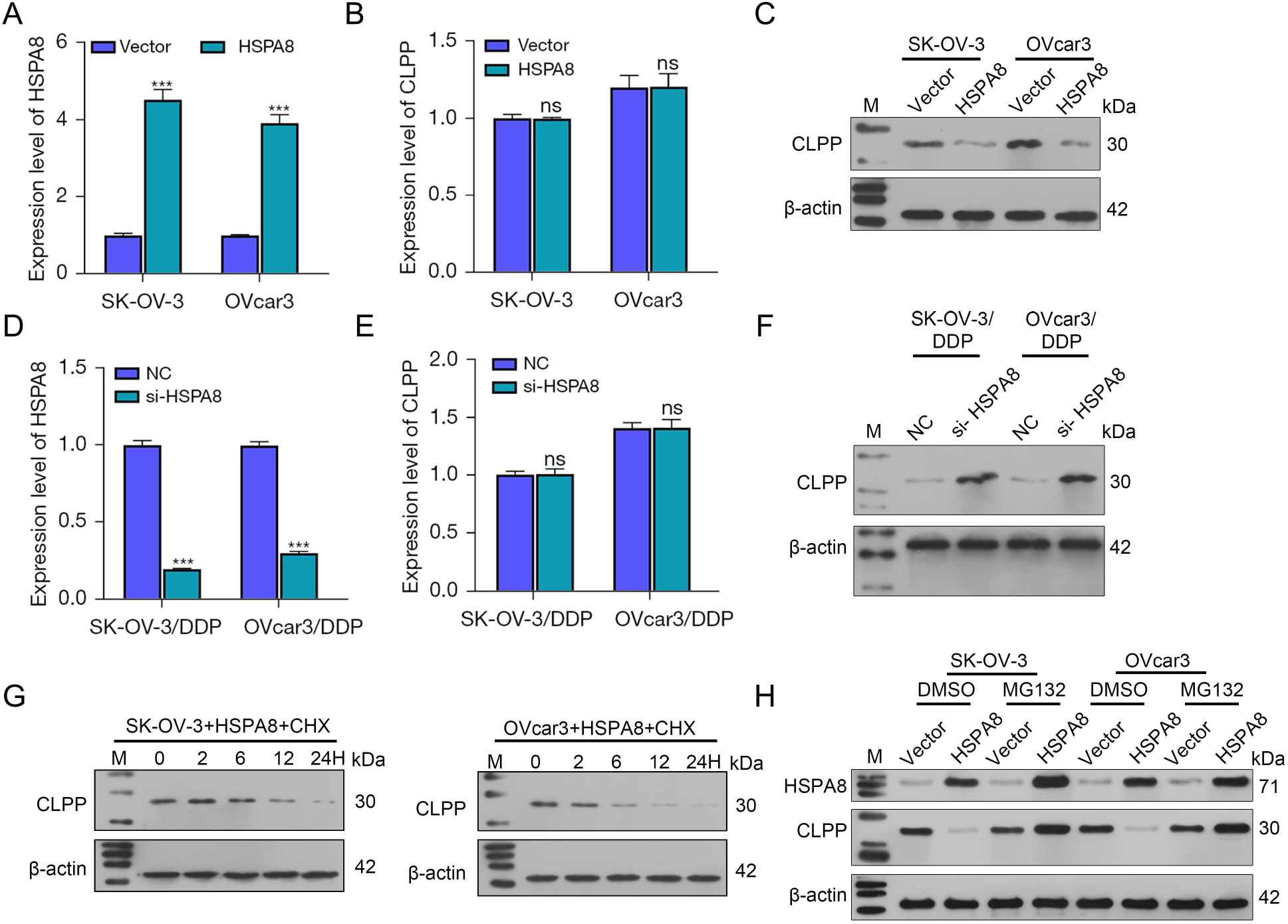
HSPA8 affected CLPP protein stability in ovarian cancer cells. (A) HSPA8 overexpression was established in the SK-OV-3 and OVcar3 cells as confirmed by real-time PCR. (B) The levels of CLPP mRNA in the transfected SK-OV-3 and OVcar3 cells were determined by real-time PCR. (C) The levels of CLPP protein in the transfected SK-OV-3 and OVcar3 cells were determined by immunoblotting. (D) HSPA8 knockdown was established in the DDP-resistant SK-OV-3 and OVcar3 cells as confirmed by real-time PCR. (E) The levels of CLPP mRNA in the transfected DDP-resistant SK-OV-3 and OVcar3 cells were determined by real-time PCR. (F) The levels of CLPP protein in the transfected DDP-resistant SK-OV-3 and OVcar3 cells were determined by immunoblotting. (G) The levels of CLPP protein in HSPA8 over-expressing SK-OV-3 and OVcar3 cells treated with CHX were determined by immunoblotting. (H) SK-OV-3 and OVcar3 cells with stable overexpression of HSPA8 were treated with a proteasome inhibitor (MG132, 20 μM). The levels of HSPA8 and CLPP proteins were detected by western blotting. ***, P < 0.001, compared to the vector or NC group. Vector, Empty carrier; NC, negative control; CLPP, caseinolytic protease P; CHX, cycloheximide; DDP, cisplatin; HSPA8, heat shock protein family A member 8; M, marker; ns, not significant; PCR, polymerase chain reaction; si, small interfering.

### HSPA8 affected wild-type ovarian cancer cell phenotypes by downregulating CLPP

As mentioned above, HSPA8 affects the stability of CLPP protein. To further investigate the dynamic effects of HSPA8 and CLPP on the DDP resistance of ovarian cancer, the wild-type SK-OV-3, and OVcar3 cells were co-transfected with HSPA8 and CLPP-overexpression plasmids. Next, the transfected cells were exposed to different concentrations of DDP (0–20 μg/mL), and the IC_50_ values were calculated based on cell viability. As shown in *Figure 5A*, HSPA8 overexpression significantly increased the IC_50_ values of the SK-OV-3 and OVcar3 cells, while CLPP overexpression partially reversed the effects induced by HSPA8 overexpression. Next, the levels of CLPP mRNA were confirmed by qRT-PCR (*Figure 5B*). Those results showed that CLPP protein levels were decreased by HSPA8 overexpression and restored by CLPP overexpression, while the levels of HSPA8 protein were not affected by CLPP overexpression (*Figure 5C*). With regard to mitochondrial autophagy and autophagy related proteins, HSPA8 overexpression increased the levels of PINK1, Parkin, and LC3-II/I proteins, while CLPP overexpression restored the levels of protein expression induced by HSPA8 overexpression (*Figure 5D*). LC3B protein expression was increased by HSPA8 overexpression and restored by CLPP overexpression (*Figure 5E*). Furthermore, cellular ROS levels and apoptosis rates were decreased by HSPA8 overexpression and restored by CLPP overexpression (*Figure 5F,5G*). These results indicated that HSPA8 could inhibit mitochondrial autophagy by down-regulating CLPP expression, and thereby promote the expression of mitochondrial autophagy-related proteins, and the interactions between LC3-II/I and mitochondrial autophagy-related proteins.

**Figure 5.**
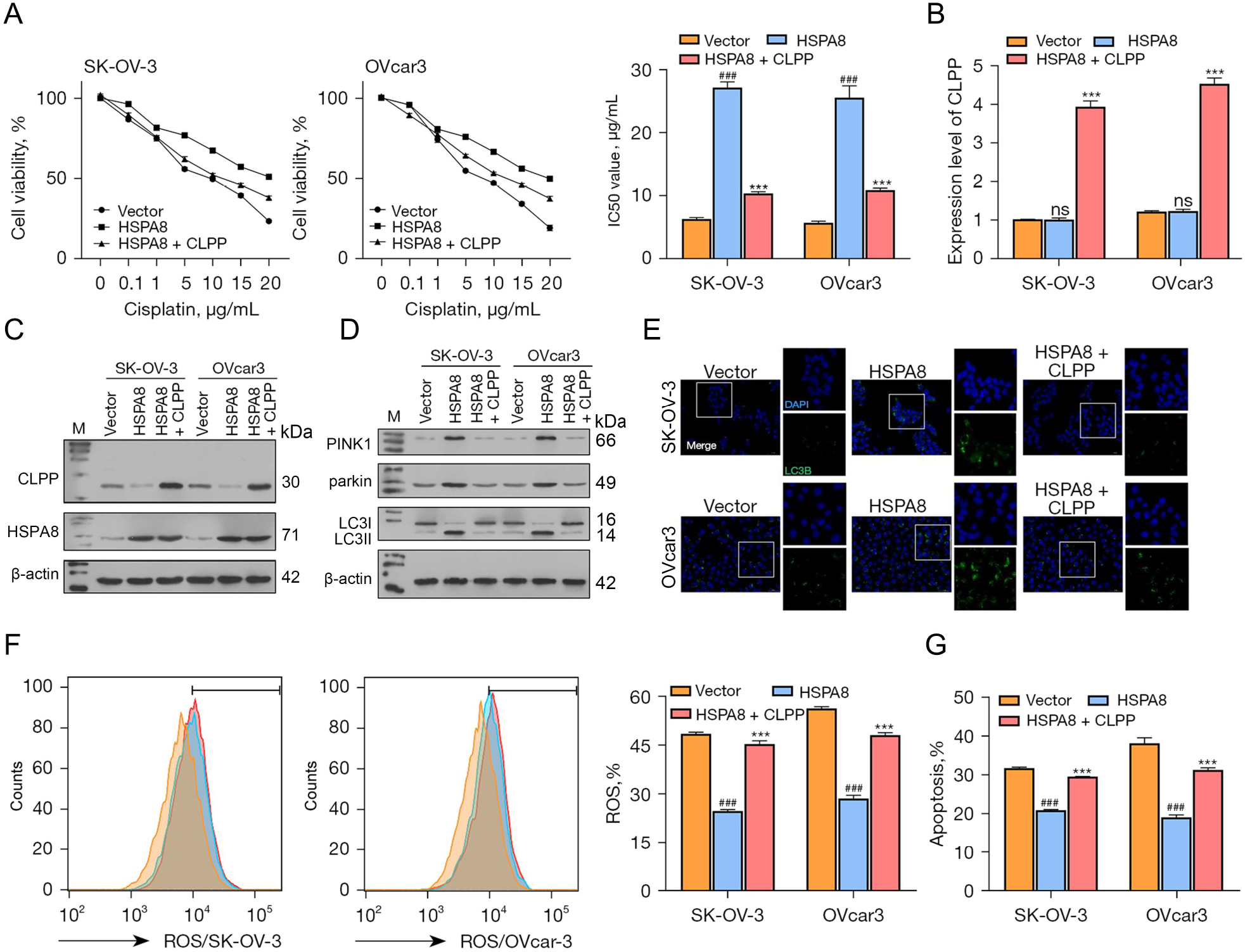
HSPA8 affected wild-type ovarian cancer cell phenotypes by downregulating CLPP. (A) Wild-type ovarian cancer cells were co-transfected with HSPA8 and CLPP overexpression plasmids and then treated with different concentrations of DDP; the IC_50_ values were calculated based on cell viability. (B) The levels of CLPP mRNA in the cells were examined by real-time PCR. (C, D) The levels of CLPP, PINK1, Parkin, and LC3II/I proteins were examined by immunoblotting. (E) The distribution of LC3B protein in the cells was examined by immunofluorescent staining (magnification x200 and x400). (F) ROS production was measured by flow cytometry. (G) Cell apoptosis was examined by flow cytometry. ^###^, P < 0.001, compared to the vector group; ***, P < 0.001, HSPA8 compared to the HSPA8 + CLPP group. CLPP, caseinolytic protease P; DDP, cisplatin; HSPA8, heat shock protein family A member 8; M, marker; ns, not significant; PCR, polymerase chain reaction; ROS, reactive oxygen species.

### HSPA8 knockdown upregulated CLPP and thus affected DDP-resistant ovarian cancer cell phenotypes

Next, the DDP-resistant SK-OV-3 and OVcar3 cells were treated with small-interfering RNA (specifically targeting HSPA8 and CLPP) to achieve the knockdown of HSPA8 and CLPP. The cells were exposed to different concentrations of DDP (0– 60 μg/mL), and the IC_50_ values were calculated based on cell viability. As shown in Figure 6A, HSPA8 interference significantly decreased the IC_50_ values of the DDP-resistant SK-OV-3 and OVcar3 cells, while CLPP knockdown partially restored the effects induced by HSPA8 knockdown. Next, the levels of CLPP mRNA were confirmed by qRT-PCR (*Figure 6B*). The levels of CLPP protein were increased by HSPA8 knockdown and restored by CLPP knockdown; the levels of HSPA8 protein were reduced by HSPA8 knockdown, but not affected by CLPP knockdown (*Figure 6C*). With regard to mitochondrial autophagy-related proteins, HSPA8 knockdown decreased the PINK1, Parkin, and LC3II/I protein levels, while CLPP knockdown restored the protein levels induced by the HSPA8 knockdown (*Figure 6D*). LC3B protein expression was decreased by HSPA8 knockdown and restored by CLPP knockdown (*Figure 6E*). The cellular ROS levels and apoptotic rates were increased by HSPA8 knockdown and restored by CLPP knockdown (*Figures 6F,6G*). Taken together, these results suggest that HSPA8 mediates cisplatin-resistant ovarian cancer cell phenotypes via CLPP dependent signaling.

**Figure 6.**
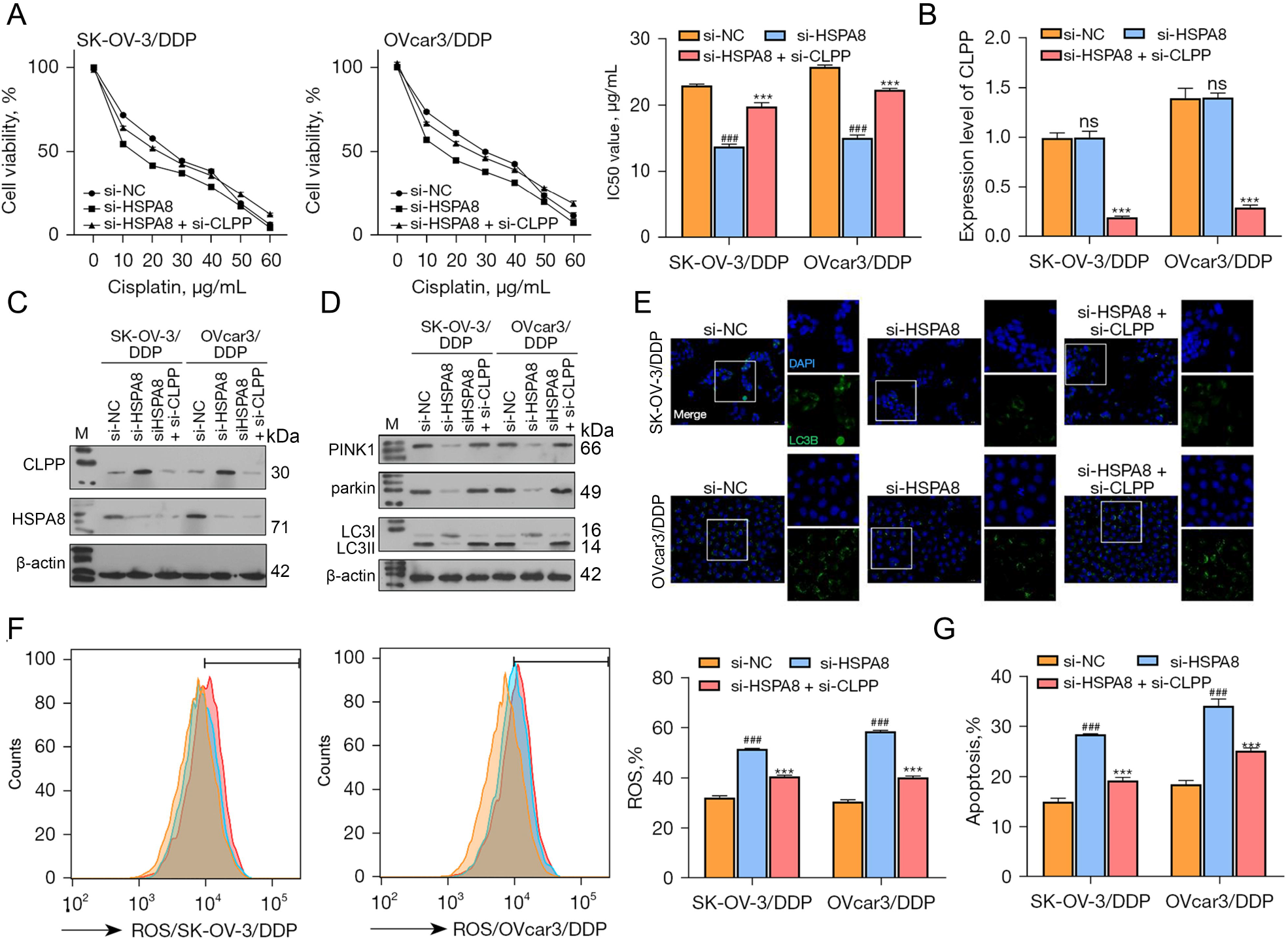
HSPA8 knockdown upregulated CLPP to affect DDP-resistant ovarian cancer cell phenotypes. (A) DDP-resistant ovarian cancer cells were transfected with HSPA8 and the CLPP interference sequence and then treated with different concentrations of DDP; the IC_50_ values were calculated based on cell viability. (B) The levels of CLPP mRNA in the cells were examined by real-time PCR. (C, D) The levels of CLPP, HSPA8, PINK1, Parkin, and LC3II/I proteins were examined by immunoblotting. (E) The distribution of LC3B protein in cells was examined by immunofluorescent staining (magnification x200× andx400). (F) ROS production was measured by flow cytometry. (G) Cell apoptosis was examined by flow cytometry. ^# # #^, P < 0.001, compared to the si-NC group; ***, P < 0.001, si-HSPA8 compared to the si-HSPA8 + si-CLPP group. si-NC, negative control; CLPP, caseinolytic protease P; DDP, cisplatin; HSPA8, heat shock protein family A member 8; M, marker; ns, not significant; PCR, polymerase chain reaction; ROS, reactive oxygen species; si, small interfering.

## Discussion

Currently, platinum agents remain the main chemotherapy drugs for ovarian cancer. DDP chemotherapy combined with surgery is also the primary treatment for malignant ovarian cancer, and a majority of seriously ill patients benefit from this therapy [34]. A platinum agent plus a taxane is the main type of chemotherapy for EOC [3]. However, the occurrence of resistance is the main reason for treatment failure. Most patients with advanced EOC suffer from disease recurrence and develop platinum resistance; after which, the choice of other therapeutic options and their effects are diminished [35]. Thus, elucidating the mechanisms of resistance occurrence and identifying drug-resistant biomarkers are vital for overcoming this dilemma and developing effective treatments.

In this study, 2 DDP-resistant ovarian cancer cell lines were established and CLPP was found to be significantly downregulated in the DDP-resistant cells. CLPP, which is a mitochondrial protease, participates in mitochondrial proteostasis and cellular stress. It is often reported to have a tumor-suppressive effect [23]. In ovarian cancer, the activation of CLPP by small molecules significantly inhibits cellular proliferation, adhesion, and metastasis, and induces G1 phase arrest, cellular stress, and cell death [27, 36, 37]. Our experiments demonstrated that knockdown of CLPP significantly increased the IC_50_ values of the wild-type ovarian cancer cells, and the IC_50_ values of the DDP-resistant ovarian cancer cells were decreased by CLPP overexpression, which is consistent with the tumor-suppressive effects of CLPP. These results provide the first evidence for involvement of CLPP in DDP resistance in ovarian cancer.

Mitochondria are the energy factories of cells, and play a key role in the metabolic reprogramming of cancers [38]. In addition to regulating bioenergetic metabolism, mitochondria also regulate key signals involved in epigenetic processes, oxidative stress, protein metabolism, and the initiation and execution of apoptosis [39]. These multiple functions give mitochondria the ability to perceive different stresses and help cells adjust to challenges from their microenvironment [39]. Mitochondrial autophagy is a selective form of autophagy which involves the PINK1-Parkin and BNIP3/NIX pathways. Those two pathways target damaged mitochondria for autophagosome degradation and reduce the levels of reactive oxygen species in mitochondria via LC3II connections, thereby promoting the survival of various types of cancer cells and tumors exposed to cytotoxic stress. LC3 occurs in two forms, designated as LC3-I and LC3-II, respectively. Both forms are produced post-translatively in various cells, and the ratio of LC3-II/I correlates with the degree to which autophagosomes are created [40]. The mammalian LC3 allotroph, LC3B, has been widely used as an autophagy membrane marker. Jin et al [42] showed that inhibition of LC3B could increase the chemical sensitivity of ovarian cancer cells. The protein kinase PINK1 and the E3 ubiquitin ligase Parkin control the specific elimination of dysfunctional or redundant mitochondria, and can thereby fine-tune the mitochondrial network and maintain energy metabolism [41]. BNIP3 is a member of the Bcl-2 protein family, which belongs to the BH3 subfamily containing only the BH3 domain [42]. The removal of damaged mitochondria is achieved via both cell autophagy and mitochondrial autophagy. NIX, a protein on the outer membrane of mitochondria, recruits LC3 II/I to form autophagosomes that wrap damaged mitochondria. This autophagy process can help to maintain intracellular stability by allowing damaged mitochondria to be cleared [43, 44]. Thus, mitochondria function is thought to be closely related to DDP resistance. Research has shown that mitochondrial alterations and abnormal mitophagy play a key role in DDP resistance [19], and inhibition of mitophagy is a promising method of intervention for combating DDP resistance in ovarian cancer [20]. In this study, we examined the effect of CLPP on mitophagy in ovarian cancer cells undergoing DDP treatment. Our data showed that CLPP knockdown promoted the expression of PINK1, Parkin, and LC3II/I proteins in the wild-type ovarian cancer cells, while CLPP overexpression decreased those levels of expression. This suggests that CLPP can regulate mitochondrial autophagy in cisplatin-resistant ovarian cancer cells.

HSPA8 is a conserved molecular chaperone that plays an indispensable role in the cellular stress response (9). High levels of HSPA8 expression have been detected in various cancer cells, and are thought to promote cancer cell proliferation and autophagy [45]. HSPA8 is highly expressed in acute myeloid leukemia, and patients with high HSPA8 expression tend to have a poor prognosis [46]. HSPA8 ablation inhibits cell proliferation and enhances the chemosensitivity of imatinib-resistant chronic myeloid leukemia cells to imatinib [47]. HSPA8 overexpression promotes cell viability and autophagy in pancreatic cancer cells [48]. In this study, our database predictions and experimental analyses showed that HSPA8 was involved in regulating CLPP protein stability. HSPA8 did not affect CLPP mRNA expression, but HSPA8 overexpression promoted the degradation of CLPP protein. In addition, we further investigated the effects of HSPA8 on ovarian cancer cells. Our data showed that HSPA8 increased the IC_50_ values and mitophagy, and inhibited ROS production and cell death. Consistent with previous studies [49, 50], our findings also suggest that HSPA8 has oncogenic functions.

## #Conclusions

In this study, we successfully generated 2 DDP-resistant ovarian cancer cell lines and found that CLPP increased the DDP resistance of ovarian cancer cells by inhibiting mitophagy and promoting cellular stress. Meanwhile, HSPA8 promotes the degradation of CLPP protein by inducing its stability. Our study explored a novel regulatory axis of DDP resistance in ovarian cancer cells and suggests promising targets (i.e., HSPA8, CLPP, and mitochondrial protein homeostasis) for overcoming DDP resistance.

## Acknowledgments

We would like to thank all those who participated in this study.

## Ethical Statement

The authors are accountable for all aspects of the work in ensuring that questions related to the accuracy or integrity of any part of the work are appropriately investigated and resolved.

## Data Availability Statement

All data are available from the corresponding author with reasonable request.

