## Supplementary material for "HSPA8-mediated stability of the CLPP protein regulates mitochondrial autophagy in cisplatin-resistant ovarian cancer cells": Table S1: Table S1.docx

**Talbe S1. Primer sequence used in PCR**

| **ID** | **Sequence(5’- 3’)** | **Product Length(bp)** |
| --- | --- | --- |
| GAPDH F | TGTTCGTCATGGGTGTGAAC | 154 |
| GAPDH R | ATGGCATGGACTGTGGTCAT |  |
| CLPP F | TGGAGCAGACGGGTCG | 125 |
| CLPP R | GAGGAGCTGTGCGATAACAA |  |
| HSPA8 F | TCAGGTTTATGAAGGCGAGC | 158 |
| HSPA8 R | TGTCCACAGCAGAGACATTG |  |
